# Antifungal activity of *Ocotea odorifera* (Vell.) Rowher, *Ocotea puberula* (Rich.) Nees and *Cinnamodendron dinisii* Schwanke essential oils

**DOI:** 10.1101/859272

**Authors:** P. Mezzomo, T.L. Sausen, N. Paroul, S.S. Roman, A.A.P. Mielniczki, R.L. Cansian

## Abstract

Biocompounds are promising tools with the potential to control pathogenic microorganisms. The medicinal plant species *Ocotea odorifera*, *Ocotea puberula* and *Cinnamodendron dinisii*, distributed along Brazilian biomes, are sources of chemical compounds of biological interest. This study aimed to evaluate the antifungal activity of the essential oils of *O. odorifera*, *O. puberula* and *C. dinisii* essential oils upon the mycotoxin producers *Alternaria alternata*, *Aspergillus flavus* and *Penicillium crustosum*. The essential oils where characterized by gas chromatography coupled to mass spectrometer (CG-MS). The majority compounds identified were: safrol (39.23%) and camphor (31.54%) in *O. odorifera*, Beta-caryophyllene (25.01%) and spathulenol (17.74%) in *O. puberula*, and bicyclogermacrene (23.19%) and spathulenol (20.21%) in *C. dinisii*. The Minimal Inhibitory Concentration (MIC) of antifungal activity considered diameters higher than 10 mm after 72 h of incubation at 30 ºC. *A. alternata* presented higher resistance to *O. odorifera* and *C. dinisii* oils. The inhibitory effect of *O. odorifera* on *A. flavus* showed stabilization at oils concentrations between 50% and 80%, increasing at 90% and 100% (pure oil) treatments. We observed that the essential oils of *O. odorifera* and *C. dinisii* have potential in the control of the analyzed fungi species. The essential oil of *O. odorifera* presented a better activity in all the assays, which can be related to the presence of safrole and phenylpropenes, compounds with known antifungal activity.

## Introduction

Chemical and synthetic biocides, largely used in conventional agriculture, are toxic to the environment and human health since their residues can contaminate and dissipate through aquatic resources (Liu et al., 2013). In contrast, natural compounds are promising tools whose do not present the contraindications or problems related to resistance development by microorganisms, as observed with conventional agrochemicals (Pinto et al., 2002). The use of biocompounds are aligned with the principle of phytotherapy, that viewing to control etiologic agents employing volatile by-products obtained from secondary metabolites of aromatic plants (Govindachari et al., 2000; Souza et al., 2012; Siqui et al., 2000; Pimentel and Burgess, 2014).

The exploration of vegetal compounds is a viable option to reduce the use of agricultural defensives (Schwan-Estrada and Stangarlin, 2005). Several extracts, resins and essential oils from plants have the potential to control pathogenic microorganisms (Gahukar, 2012; Garcia et al., 2008; Küster et al., 2009). The extraction of plant products evolves simple and low cost methods whose remits to effective potential to the technological development, employment in industrial plants and home use as well.

The use of native species as sources of organic composts can be a strategy to promote the sustainable manage of natural resources allied to economic interests (Vedovatto et al. 2015). *Ocotea odorifera* (Vell.) Rowher and *Ocotea puberula* (Rich.) Nees are native tree species from Brazil, found in Araucaria Forest with phytophysiognomy of Mixed Ombrophilous Forest (Lorenzi, 2008). These species present wide geographic distribution in South America and have ecological importance in the recovering of degraded areas (Zangaro et al., 2003; Montrucchio et al., 2012).

The phytochemical metabolism of *O. odorifera* and *O. puberula* includes the synthesis of flavonoids as kaempferol and quercetin, steroids and sesquiterpenes (Costa, 2000; Lordello et al., 2000). The major compounds of their essential oil are safrole and Beta-caryophyllene, which are used in pharmaceutical industries for drug production due to sudorific, antirheumatic, antiseptic, diuretic and repellent properties (Lorenzi, 2016; Pinto Junior et al., 2010).

*Cinnamodendron dinisii* Schwanke is a pioneer specie of Atlantic Forest also encountered in regions of Mixed Ombrophilous features (Souza and Lorenzi, 2012), with geographical distribution between Brazilian States of South and Southeast regions (Lorenzi, 2016). These species are sources of chemical compounds of biological interest. The chemical profile of *C. dinisii* indicates the presence of drimane sesquiterpenes, a class of hydrocarbons with documented bactericidal, antifungal, cytotoxic, phytotoxic and piscicide effects (Jansen and Groot, 2004).

*Alternaria alternata*, *Aspergillus flavus* and *Penicillium crustosum* are fungal species that produces mycotoxins, secondary metabolites that can present toxicity to humans and other animals (Bennett and Klich, 2003) by food products contamination. Mycotoxins can induce carcinogenic, immunotoxic, hepatotoxic and nephrotoxic responses in the exposed organisms (Bennett and Klich, 2003). Approximately 300 compounds are recognized as mycotoxins and about 30 of them are classified as dangerous to the human health. According to Streit et al. (2013), more than 70% of the samples from raw material utilized to animal food are positive to at least one type of mycotoxin.

Considering the need to develop sustainable alternatives to control phytopathogenic organisms, in this study, we develop an original research to evaluate the antifungal activity of the essential oils of *O. odorifera*, *O. puberula* and *C. dinisii* upon the mycotoxin producers *A. alternata*, *A. flavus* and *P. crustosum*.

## Materials and methods

Samples of *O. odorifera*, *O. puberula* and *C. dinisii* were collected in the year of 2015 at different locations of the municipality of Erechim, southern Brazil. Approximately 3kg of fresh leaves of each specie were collect, weighted and dried in air heater at 30 °C, until reach constant weight. Then, leaves where triturated until converted to a homogeneous powder. The essential oils were obtained by hydrodistillation in Clevenger apparatus and the final product were stored in glass recipients at −20 °C. Each sample was hydrodistilled using the same technique (Clevenger), and each of the essential oils were analyzed by GC-MS under the same conditions.

The chemical composition of the essential oils was analyzed using a Shimadzu QP 5050A series gas chromatograph coupled to mass spectrometer (GC-MS), with a DB-5 fused silica capillary column (30 m × 0.25 mm internal diameter ×0.25 µm film thickness). Samples of 50.000 ppm, diluted in hexane, were applied on chromatographic column. The injection port was in split 1:20, with injector temperature at 250 °C and interface temperature at 250 °C, in DB5 column, with flow of 1 mL/min, 1.6 kV detector and solvent cutting at 3.5 min. The temperature program was the following: initial temperature of 50 °C, during 3 min, and ramped at 5 °C/min until 130 °C, then 15 °C/min until reach 210 °C/min by 7 minutes and 20 °C/min until reach 250 °C by 10 minutes. The compounds were identified by comparison with the mass spectrum library (The Wiley Registry of Mass Spectral Data, 7th ed.) and the Kovats index (Adams, 2007).

The antifungal activity was evaluated using different oil concentrations, from pure oil (100%, without additives) to dilutions of 10, 20, 30, 40, 50, 60, 70, 80 and 90% of oils in Tween 80 (Synth, BR) and distilled sterile water. The strains of *A*. *alternata*, *A*. *flavus* and *P*. *crustosum* were obtained from Agricultural Research Service (ARS Culture Collection - NRRL) and stored in PDA medium (Potato Dextrose Agar) at 4 °C. The toxicity assays were made by diffusion method in solid medium (Hadaceck and Granger 2000). In this assay, 1 ml of fungal suspension containing around 106 CFU mL-1 was mixed with PDA medium, previously melted (40−50 °C), and distributed in petri dishes. After solidification, sterile glass cannulates were used to make cavities of 6 mm diameter which, where were deposited the test oil solutions (50 µl per cavity) and the negative and positive controls that were, respectively, Tween 80 and commercial Ketoconazole (Ibasa, BR). The plates were then incubated at 30 °C for 72 hours, with standard observations in intervals of 24 hours. After the incubation time, the halos diameters were measured and the Minimal Inhibitory Concentration (MIC) is defined as the concentration able to inhibit the fungal growth at halos higher than 10 mm diameter. All experiments were conducted in triplicate and the results are expressed as mean ± standard deviation. The data were analyzed by one-way ANOVA plus Tukey-test, with *p* <0.05 being considered significant. Statistical analyses were performed only when the inhibitory halo reached the MIC, as defined above (at least 10 mm diameter).

## Results and Discussion

The essential oils of *O. odorifera*, *O. puberula* and *C. dinisii* yielded a mean of 0.96, 0.51 and 0.62 mL of oil per 100 grams of dried leaves, respectively, after 2 hours of extraction. The chromatography analyses identified 8 majority compounds in *O. odorifera*, 14 in *O. puberula* and 15 in *C. dinisii* (Table 1).

**TABLE 1.** Majority chemical composition of *O. odorifera*, *O. puberula* and *C. dinisii* essential oils.

| # | Compound name | RI (Kovats) | *O. odorifera* (%) | *O. puberula* (%) | *C. dinisii* (%) |
| --- | --- | --- | --- | --- | --- |
| 1 | Alpha-thujene | 0909 | - | 3.71 | - |
| 2 | Alpha-pinene | 0945 | 2.84 | 4.53 | - |
| 3 | Camphene | 0953 | 5.05 | - | - |
| 4 | Beta-pinene | 0980 | 1.23 | 7.21 | - |
| 5 | Limonene | 1029 | - | 3.50 | 1.03 |
| 6 | Sabinene | 1068 | 2.52 | - | - |
| 7 | Camphor | 1143 | 31.54 | - | 2.73 |
| 8 | Alpha-terpineol | 1148 | - | - | 5.34 |
| 9 | Amyl acetate | 1275 | - | - | 2.72 |
| 10 | Safrol | 1285 | 39.23 | - | - |
| 11 | Terpinyl acetate | 1352 | - | - | 11.31 |
| 12 | Geranyl acetate | 1382 | - | - | 1.25 |
| 13 | Calarene | 1385 | - | 3.61 | 2.48 |
| 14 | Beta-elemene | 1393 | - | 9.72 | - |
| 15 | Isocaryophyllene | 1438 | - | - | 2.72 |
| 16 | Alpha-humulene | 1457 | - | 2.23 | 1.83 |
| 17 | Beta-caryophyllene | 1475 | - | 25.01 | 1.34 |
| 18 | Germacrene D | 1485 | - | 3.10 | 1.14 |
| 19 | Bicyclogermacrene | 1510 | - | 9.31 | 23.19 |
| 20 | δ-Cadinene | 1523 | - | 3.12 | 2.8 |
| 21 | *trans*-Nerolidol | 1552 | - | 4.41 | 1.49 |
| 22 | Spathulenol | 1576 | 3.87 | 17.74 | 20.21 |
| 23 | *trans*-Muurolol | 1682 | - | 3.79 | - |
| 24 | Alpha-farnesol | 1697 | 2.35 | - | - |
|  | Monoterpene hydrocarbons |  | 11.64 | 18.95 | 1.03 |
|  | Oxygenated monoterpenes |  | 31.54 | - | 23.35 |
|  | Phenylpropene |  | 39.23 | - | - |
|  | Sesquiterpene hydrocarbons |  | - | 56.1 | 35.5 |
|  | Oxygenated sesquiterpenes |  | 6.22 | 25.94 | 21.7 |
|  | **Total** |  | **88.63** | **100** | **81.58** |

*O. odorifera* essential oil was characterized mainly by volatile components (oxygenated and non-oxygenated monoterpenes and phenylpropenes), that accounted about 82% of the total oil composition, with safrole (39.23%) and camphor (31.54%) representing 70% of the identified compounds.

For the *O. puberula* essential oils, there was a predominance of oxygenated and non-oxygenated sesquiterpenes, with Beta-caryophyllene (25.01%) and spathulenol (17.74%) being the two majoritarian compounds. The oil of *C. dinisii* presented 24% of oxygenated and non-oxygenated monoterpenes and 57% of oxygenated and non-oxygenated sesquiterpenes, essentially bicyclogermacrene (23.19%) and spathulenol (20.21%). The chemical composition of the major compounds founded in the three essential oils of the analyzed species is described in the Table 1.

Several studies have demonstrated antifungal and antimicrobial properties of plants essential oils (Oxenham et al., 2005), that, essentially, are produced as a defense against pathogenic organisms (Ahmet et al., 2005). Specific responses of phytopathogens to the plant compounds can be found in the literature, as well as the chemical classes that are most effective in the control of fungal and bacterial species (Tewarri and Nayak, 1991; Amadioha, 2000; Okigbo and Nmeka, 2005).

The antifungal analyses indicated inhibitory potential of the essential oils of *O. odorifera* and *C. dinisii* had upon *A. alternata*, *A. flavus* and *P. crustosum* in concentrations higher than 50% of the initial dilution (Table 2). However, the oil of *O. puberula* did not present satisfactory inhibition, with halos lower than 10 mm diameter in all the tested concentrations, indicating low biological activity of this specie in relation to the target organisms, according to the established MIC.

**TABLE 2.** Antifungal activity of *O. odorifera*, *O. puberula* and *C. dinisii* essential oils upon *A. alternata*, *A. flavus* and *P. crustosum*.

| *Ocotea odorifera* | Halo diameter (mm)* |  |  |
| --- | --- | --- | --- |
|  | *A. alternata* | *A. flavus* | *P. crustosum* |
| 50% | 9.90<sup>c</sup> ± 0.46 | 10.93<sup>d</sup> ± 1.00 | 9.07<sup>e</sup> ± 0.51 |
| 60% | 11.13<sup>c</sup> ± 0.55 | 12.07<sup>cd</sup> ± 0.35 | 10.97<sup>d</sup> ± 0.95 |
| 70% | 12.63<sup>b</sup> ± 0.55 | 13.40<sup>bc</sup> ± 0.36 | 12.03<sup>cd</sup> ± 0.15 |
| 80% | 13.13<sup>b</sup> ± 0.38 | 14.47<sup>ab</sup> ± 0.35 | 13.80<sup>bc</sup> ± 0.75 |
| 90% | 13.60<sup>ab</sup> ± 0.44 | 14.97<sup>a</sup> ± 0.35 | 15.37<sup>ab</sup> ± 0.60 |
| 100% | 14.83<sup>a</sup> ± 0.47 | 15.77<sup>a</sup> ± 0.49 | 16.57<sup>a</sup> ± 0.70 |

| *Ocotea puberula* | Halo diameter (mm)* |  |  |
| --- | --- | --- | --- |
|  | *A. alternata* | *A. flavus* | *P. crustosum* |
| 50% | 0.92<sup>c</sup> ± 0.61 | 0.50<sup>d</sup> ± 0.29 | 0.93<sup>b</sup> ± 0.26 |
| 60% | 0.50<sup>c</sup> ± 0.42 | 0.73<sup>d</sup> ± 0.16 | 1.53<sup>b</sup> ± 0.45 |
| 70% | 0.40<sup>c</sup> ± 0.12 | 1.10<sup>cd</sup> ± 0.29 | 2.25<sup>b</sup> ± 1.06 |
| 80% | 0.97<sup>c</sup> ± 0.07 | 2.10<sup>c</sup> ± 0.10 | 5.23<sup>a</sup> ± 0.87 |
| 90% | 2.73<sup>b</sup> ± 0.40 | 3.79<sup>b</sup> ± 0.36 | 5.37<sup>a</sup> ± 1.12 |
| 100% | 4.30<sup>a</sup> ± 0.72 | 5.37<sup>a</sup> ± 0.93 | 7.45<sup>a</sup> ± 1.40 |

| *Cinnamodendron dinisii* | Halo diameter (mm)* |  |  |
| --- | --- | --- | --- |
|  | *A. alternata* | *A. flavus* | *P. crustosum* |
| 50% | 6.40<sup>c</sup> ± 1.14 | 6.38<sup>c</sup> ± 0.63 | 15.11<sup>a</sup> ± 6.11 |
| 60% | 8.24<sup>bc</sup> ± 1.05 | 6.40<sup>c</sup> ± 0.62 | 18.14<sup>a</sup> ± 7.42 |
| 70% | 9.10<sup>b</sup> ± 1.40 | 6.93<sup>c</sup> ± 2.91 | 19.33<sup>a</sup> ± 4.50 |
| 80% | 10.47<sup>b</sup> ± 0.57 | 7.70<sup>c</sup> ± 1.21 | 19.33<sup>a</sup> ± 8.08 |
| 90% | 16.17<sup>a</sup> ± 0.77 | 15.00<sup>b</sup> ± 2.00 | 24.48<sup>a</sup> ± 3.64 |
| 100% | 16.07<sup>a</sup> ± 0.15 | 20.20<sup>a</sup> ± 1.81 | 30.00<sup>a</sup> ± 5.57 |

\* Mean ± standard derivation following by same letter in the columns not presented statistical differences ( $p < 0.05$ ) in accordance with Tukey test.

The search for natural products able to be used in biological control of pathogens is a trend towards sustainability. Aromatic and medicinal plants became popular in the field of disease control, since their essential oils usually contain antimicrobial properties due to their spectrum of secondary metabolites such as alkaloids, quinones, flavonoids, glycosides, tannins and terpenoids (Balakumar et al. 2011; Gillitzer et al. 2012) that, isolated or synergistically, may exhibit such of these properties.

The two major compounds of the essential oil of *O. Odorifera* that were identified in this work are known for their commercial interest. Safrol is used as raw material in the manufacture of piperonal (or heliotropin), an organic compound used in fragrances and flavors; and also, in the synthesis of some synthetic agents used in the formulation of pyrethroid insecticides (Keil, 2007). Camphor was previously described by its antifungal potential, especially on *A. flavus*, which is in accordance with the data obtained in this study (Rasoli and Owlia, 2005; Kumar et al., 2007; Rasoli et al., 2008; Bluma et al., 2008; Adjou et al., 2012;).

Regarding the essential oil of *O. odorifera*, the antifungal profile was similar for the three target fungi species (*A. alternata*, *A. flavus* and *P. crustosum*), with the inhibition halo generated by the concentration of 90% being statistically equivalent to the concentration of 100% (pure oil). In all cases, a good linear correlation was observed between oil concentration and antifungal performance (Figures 1A-C).

**FIGURE 1.**
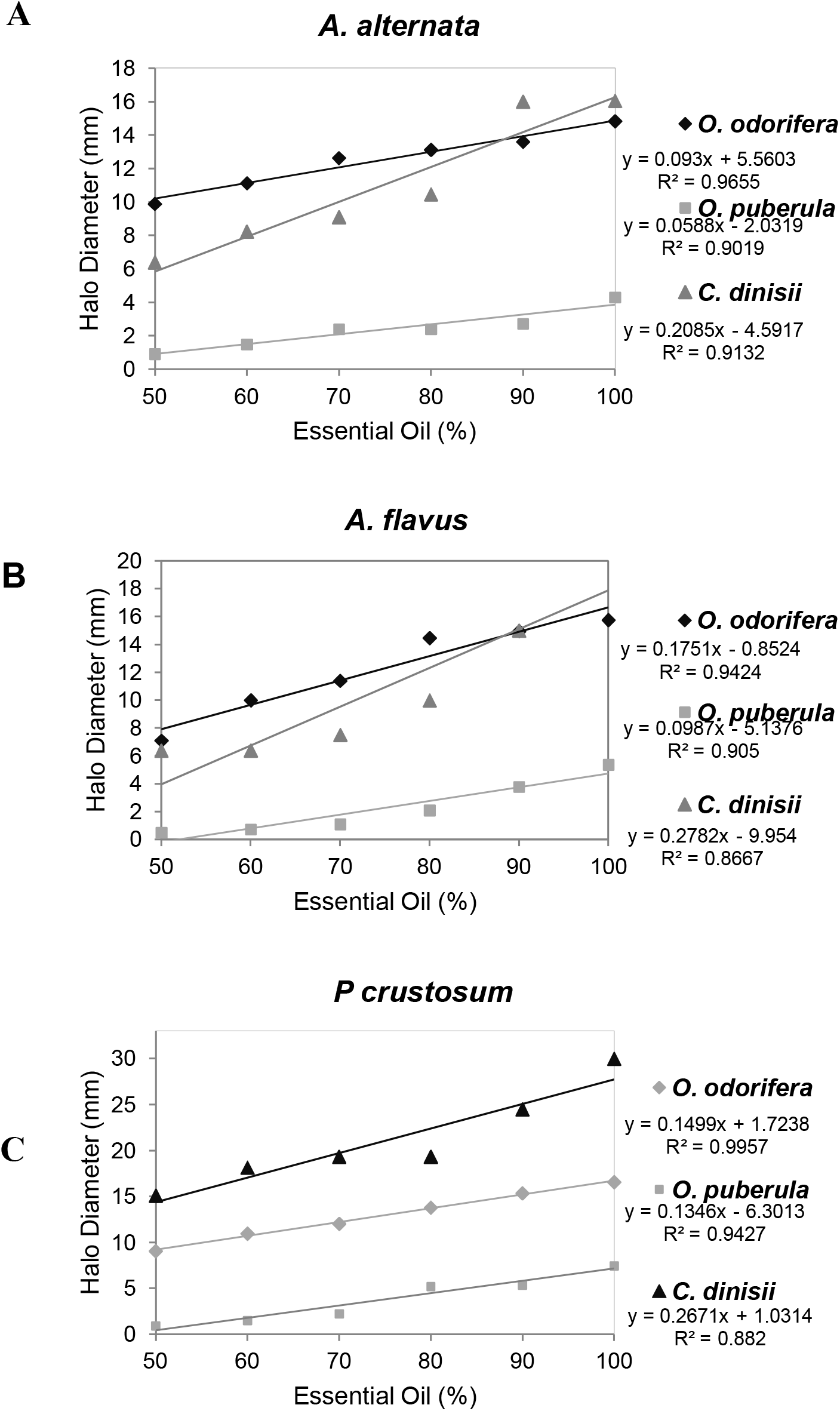
Correlation between essential oil concentrations and halo of growth inhibition of *A. alternata*, *A. flavus* and *P. crustosum*. (A) *O. odorifera* oil, (B) *O. puberula* oil and (C) *C. dinisii* oil.

Beta-caryophyllene, the predominant component of *O. puberula*, can acts directly in fungal inhibition and have antimicrobial, anticancer and anti-inflammatory properties (Cantrell et al., 2005; Asdadi et al., 2015). Sesquiterpene hydrocarbons as spathulenol and bicyclogermacrene, identified in *O. puberula* (spathulenol 17.74%) as well in *C. dinisii* (spathulenol 20.21% and bicyclogermacrene 23.19%) present antimicrobial and moderately cytotoxic activity (Limberger et al., 2004, Constantin et al., 2001; Cysne et al., 2005). However, in this work *O. puberula* showed weak antifungal inhibition, indicating that the chemical composition of their oil is not sufficiently efficient against *A. alternate*, *A. flavus* and *P. crustosum* species.

The *C. dinisii* oil have the higher efficiency against *P. crustosum*, reaching an inhibition halo of 30 mm diameter at the concentration of 100%, which had equal statistical significance in relation to the other tested concentrations (from 50 up to 90%). The effect upon *A. alternata* and *A. flavus* reached top values in the presence of 90% (16 mm halo) and 100% (20 mm halo) of oil concentration, respectively. The halos generated by *O. odorifera* in *A. flavus* also showed stabilization in essential oil concentration between 70% and 80% (Table 2 and Figure 1B).

The correlation between the essential oils of *O. odorifera* and *C. dinisii* and the halos diameter generated for *A. alternata* presented a stabilization of the biological activity in the concentration of 90% of oil (Figure 1A). This result, associated with the halo formation only in concentrations above 50%, is an indicative of a higher resistance of *A. alternata* to these two essential oils. For *A. flavus*, was observed a stabilization in the halo generated by *C. dinisii* oil with concentrations between 50% and 80% (Table 2 and Figure 1), with increase in essential oil halos diameter in 90% and 100%.

The mechanism through which compounds from secondary metabolism of plants inhibit the growth of pathogenic organisms, such as fungi, are not yet clearly established. However, studies have reported that some plant essential oils are responsible for DNA damage in animals and microorganisms, presenting genotoxic and mutagenic potential, suggesting that its use should be carefully evaluated and evidences the need for new research to understand the effects of these essential oils on different organisms (Vilar et al. 2008). Despite the lack of a complex understanding of the mechanisms itself, the contribution of biocompounds to the biological control is widely evidenced in the literature and are promising for its practical application in biological control of phytopathogens, such as fungi.

## Conclusions

The essential oils from *O. odorifera* and *C. dinisii* have inhibitory effect upon the growth of *A. alternata*, *A. flavus* and *P. crustosum*, indicating their potential in the control of this phytopathogens. The phytochemical characterization of native tree species has social, economic and ecologic impact, once the conservation of forestry natural resources can be improved by the development of alternatives for their sustainable use. In accordance with this concept, the results of this work indicated the potential of two plant species from Brazilian ecosystems as natural biocides against economic relevant fungi pathogens.

## Acknowledgements

The authors thanks CNPq, FAPERGS and CAPES for the financial support and scholarships provided.

